# tUbe net: a generalisable deep learning tool for 3D vessel segmentation

**DOI:** 10.1101/2023.07.24.550334

**Authors:** Natalie A. Holroyd, Zhongwang Li, Claire Walsh, Emmeline Browng, Rebecca J. Shipley, Simon Walker-Samuel

**Affiliations:** Centre for Computational Medicine Division of Medicine University College London; Department of Mechanical Engineering University College London

**Keywords:** deep learning vasculature segmentation

## Abstract

Deep learning has become an invaluable tool for bioimage analysis but, while open-source cell annotation software such as cellpose are widely used, an equivalent tool for three-dimensional (3D) vascular annotation does not exist. With the vascular system being directly impacted by a broad range of diseases, there is significant medical interest in quantitative analysis for vascular imaging. We present a new deep learning model, coupled with a human-in-the-loop training approach, for segmentation of vasculature that is generalisable across tissues, modalities, scales and pathologies. To create a generalisable model, a 3D convolutional neural network was trained using curated data from modalities including optical imaging, computational tomography and photoacoustic imaging. Through this varied training set, the model was forced to learn common features of vessels cross-modality and scale. Following this, the pre-trained ‘foundation’ model was fine-tuned to different applications with a minimal amount of manually labelled ground truth data. It was found that the foundation model could be specialised to a new datasets using as little as 0.3% of the volume of said dataset for fine-tuning. The fine-tuned model was able to segment 3D vasculature with a high level of accuracy (DICE coefficient between 0.89 and 0.98) across a range of applications. These results show a general model trained on a highly varied data catalogue can be specialised to new applications with minimal human input. This model and training approach enables users to produce accurate segmentations of 3D vascular networks without the need to label large amounts of training data.

## 1. Introduction

A broad range of diseases directly involve the vascular system, and as such the organisation and structure of blood vessels is of inherent medical interest. For example, blood vessel networks become highly disrupted following angiogenesis in cancer (1), inflammation in response to infection induces changes in vascular structure (2), and blockages causing loss of perfusion has serious consequences in cardiac infarction (3), stroke (4) and diabetes (5). Many bioimaging techniques have been developed to characterise both normal and diseased blood vessel networks, across a range of length scales, organs and pathologies. However, the ability to quantify and fully characterise the complex, hierarchical nature of blood vessels still presents a significant challenge, particularly when consideration of their three-dimensional structure is required.

Deep learning (DL) with convolutional neural networks has rapidly become the state of the art approach for semantic segmentation (labelling each pixel according to a predefined set of classes) in medical imaging due its accuracy and generalisability. However, it usually requires a large and varied array of training data, complete with paired ground truth labels. Whilst image data is generally widely available, acquiring manually-defined labels is a substantial and often challenging undertaking and introduces a significant bottleneck to implementation. Accordingly, most DL models to date have been trained using a limited set of data from a single pathology or organ, imaged with a single modality, at a single length scale. In particular, much of the field is focused on segmenting retinal vasculature from two-dimensional colour fundus images (6; 7; 8; 9; 10). Models have also been created for segmenting brain vasculature in three dimensions (11; 12). Typically, each new application requires a new model, trained on new data.

In the field of cellular imaging, cellpose (13) and cellpose2 (14) have become highly popular open-source tools that employ an alternative, a human-in-the-loop workflow: where a general model is trained most of the way and then fine-tuned on a small amount of the end-users’ data in order to specialise the model to their needs. This approach minimises manual labelling and increases the accessibility of deep learning tools for users without the computational resources or sufficient labelled data to train a model from scratch. Similar approaches are also available for vessel segmentation in the form of proprietary software (for example, the Imaris 10.0 Filament Tracer and the Amira-Avizo XFibre extension) but no open-source tools exist for this purpose.

In this study, we aimed to leverage the ubiquitous tube-like structuring of blood vessels to create a model that can be used to segment three-dimensional image data containing blood vessels, acquired with multiple modalities, and from multiple organs. This is achieved by implementing a training strategy that prioritises a highly varied multi-modal training set over volume of data. Our carefully curate This way dataset, we develop a generalisable vessel-finding model with far less pre-training data than typically reported, and demonstrate that it can be fine-tuned to new tasks with a minimal amount of user-labelled data.

### 1.1. Semantic Segmentation With U-Nets

Convolutional neural networks (CNN) consist of a sequence of convolutional layers (a deep network) with weights that can be trained to enable it to detect image features at multiple length scales. U-Nets are a class of CNN architecture consisting of encoding layers that feed into a symmetric set of decoding layers (15). In the encoding arm, alternating convolutional and pooling layers downsample the data and convert the input data into a latent representation. Once encoded, the decoding arm converts the latent representation into a new image space, by sequential upsampling using deconvolutional (or transpose convolutional) layers. The downsampling and upsampling branches are symmetric, giving a ‘U’ shaped architecture. Skip connections enable direct communication between corresponding encoding and decoding layers, primarily to help overcome the vanishing gradient problem that affects very deep CNNs and has been found to improve the accuracy of segmentation of biological images (16).

#### 1.2. Bottlenecks In ML-based Segmentation

Collating sufficient data to accurately train a deep neural network is one of the major challenges in deep learning. In the field of medical imaging there is generally no shortage of raw data, but the corresponding ground truth labels are more challenging to obtain. Images must be manually segmented to create paired datasets, which is highly labour-intensive and potentially error-prone (10). The training data must also comprise a representative sample of the range of the data the model will be expected to classify upon application, to enable sufficient generalisability (17).

Multiple approaches have been developed to overcome these challenges. In addition to creating architectures that require less training data, such as the U-Net, researchers have attempted to make smaller datasets go further using data augmentation (18; 19), and have employed pre-training (or transfer learning) strategies (20; 21; 22). Pre-training leverages the ability of CNNs to detect features which are characteristic of a broad range of image domains; as such, a CNN trained on one domain can be transferred to another domain using less data than would be required to train it from scratch. Similarly, synthetic data has been used to address training bottlenecks: models have been trained using either purely synthetic data (23; 24), or a combination of real and synthetic data (25; 26), for which the ‘ground truth’ labels are inherently known. However, the extent to which models trained on synthetic data may be meaningfully applied to physiological data remains an open question (27).

### 1.3. Semantic Segmentation Of Blood Vessels

CNNs have been successfully used to label blood vessels in two-dimensional (2D) images and have been widely applied in the segmentation of retinal vasculature from image data (6; 7; 8; 9; 10). This is aided by the fact that several hand-labelled data sets are available to allow developers to benchmark new models and approaches (e.g. DRIVE (28), Structured Analysis of the Retina (STARE) (29) and FIVES (30)).

Segmentation of three-dimensional (3D) vasculature, on the other hand, presents a greater challenge due in part to the much higher demands of manual segmentation, higher computational expense, and the lack of ready-labelled libraries against which to benchmark. Furthermore, much of the research in this area has been limited to brain vasculature; for example, Ref. 11 used a 3D CNN to segment the vascular network from dual-stained mouse brains, imaged using light-sheet microscopy, in their VesSAP pipeline. Likewise, DeepVesselNet (12) was trained with two sets of brain data: clinical MRA and rat brain micro-CT. There are few examples of applications outside of the brain, for example Ref. 31, which demonstrated the effectiveness of an integrated U-net and graphical neural network on two clinical contract-enhanced liver CT datasets. To our knowledge, no work has been published in which DL is used to segment 3D microvasculature outside of the brain.

While these examples point to many potential applications for 3D CNN image analysis, both in research and in the clinic, there is clearly a need for models that can be applied to a wider range of tissues and data sources. In particular, there is a need for models that can segment vasculature at the micron scale in organs other than the brain and retina. Here, we explore the use of transfer learning to address bottlenecks in blood vessel segmentation, and aim to create a ‘generalised’ 3D vessel segmentation model that can be fine-tuned to multiple applications.

## 2. Methods

### 2.1. Model Architecture

A convolutional neural network was produced in tensorflow version 2.3.0 by adapting the U-Net architecture for 3D data. The model consists of six convolutional layers followed by five deconvolutional layers. The convolutional layers (purple, Fig. 1) consist of two 3D convolutions (kernel size 3 × 3 × 3) followed by a Leaky ReLU activation function (alpha = 0.2) and group normalisation. A pooling function follows each layer, downsampling the data by a factor of two in each dimension - effectively halving the resolution. Deconvolutional layers (orange) increase the resolution of the data using a deconvolving function (‘conv3DTranspose’), followed by two standard convolutions, again with a Leaky ReLU activation function. The output layer is a convolution with a single filter and a softmax activation function to give a pseudo-probabilistic output for each pixel belonging to each class (i.e. inside or outside a vessel). Concatenations are used to ‘skip’ layers by transferring feature maps directly from the encoding branch to the symmetric layer of the decoding branch, which has been shown to improve convergence speed in deep networks (16). A 30 % dropout is included after each layer to reduce the risk of overfitting to the training data.

**Figure 1:**
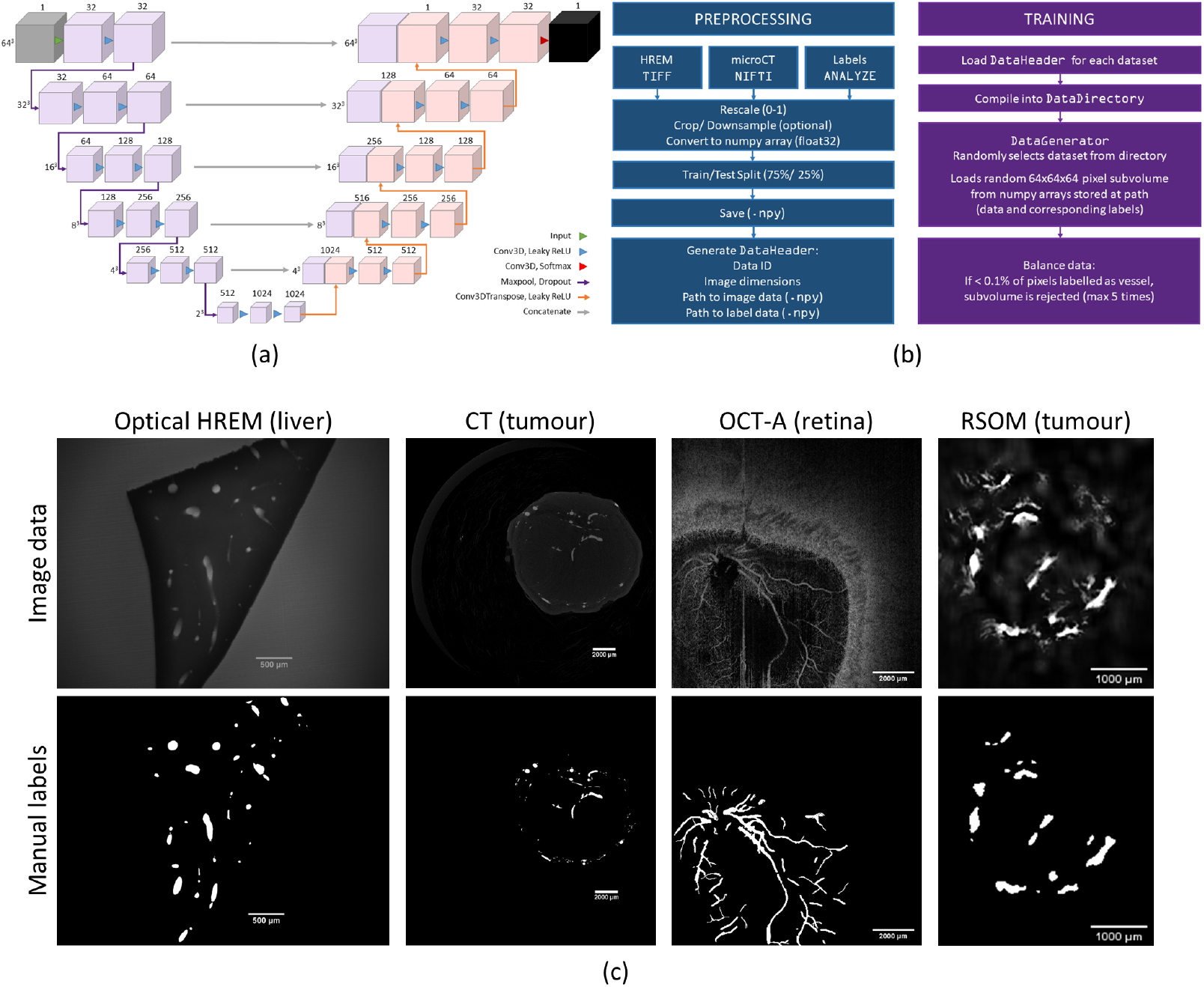
(a) This schematic shows the architecture of the tubenet model: the purple and blue boxes represent multi-channel feature maps, with the number of channels denoted by the number above the box. The dimensions to the side of each layer give the x-y-z dimensions of the feature maps in that layer. The network takes an image (grey) as an input and gives a map of probabilities as an output (black). The arrows represent operations, as summarised in the image key. (b) An overview of the data pre-processing pipeline, and method of loading data sub-volumes from the disk storage during training. The use of .npy file format and method of reading in data allows the model to train on larger datasets that cannot fit within the computer’s RAM. (c) Example images taken from the three-dimensional image data used in training. 3D vessel images from four different modalities, and three different tissue types, were manually segmented to create a varied training set. Here, a 2D slice is shown from each dataset (top row), along with the corresponding manual labels (bottom row).

### 2.2. Attention Layers

A second version of the model was created in which skip connections were replaced with attention layers. Attention mechanisms are used to capture correlations in the data across scales or channels. Here, an attention map was computed between feature maps of the same resolution in the encoding and decoding branches, similar to the method proposed in Ref. 32, but using the Luong model of attention (33) rather than additive attention. The attention map for a given layer of the U-net is produced by first transforming both the encoding branch feature map (*q*) and decoding branch feature map (*k*) via a convolution with a trainable 3×3×3 kernel. The resulting matrices are combined and then passed through another convolution layer with a 1 × 1 × 1 kernel and sigmoid activation to produce the final attention map. The attention map was then multiplied element-wise with the decoding branch feature map.

### 2.3. Model Hyperparameters

Model hyperparameters were determined using a random search function (15 iterations) to compare a range of initial learning rates (0.00001, 0.0001, 0.001), activation functions (ReLU, Leaky ReLU alpha=0.1, Leaky ReLU alpha=0.2, Leaky ReLU alpha=0.3), and dropout rates (10%, 20%, 30%). Evaluating the model performance on hold-out data (see section 2.5), the optimal combination of hyperparameters was found to be an initial learning rate of 0.001, leaky ReLU activation function with alpha=0.2, and a dropout rate of 30%.

### 2.4. Correcting For Class Imbalance

In classification problems where one class is more common than the other, ML algorithms can become biased towards the better represented class. This is common in image data containing blood vessels, as blood volume in biological tissue typically 3 % to 5 %. To counter this, the training data generator was weighted towards batches of image subvolumes containing at least 0.1% vessels. This method meant that the model would still see some batches containing no vessels, but far less often than would otherwise be the case.

Three loss functions designed to handle imbalanced classes were compared: weighted categorical cross entropy (WCCE), DICE binary cross entropy (DICE BCE), and focal loss. WCCE used pixel-wise cross entropy, a measure of similarity between distributions, commonly used for segmentation and classification tasks, and scales the contribution of each pixel based on the ground truth class. Class weights were assigned in proportion to the inverse of the class size. DICE-BCE is a sum of the DICE coefficient and Binary Cross Entropy (BCE). The DICE coefficient is robust to imbalanced classes as it takes into account recall and precision, while BCE provides a smooth gradient for backpropagation (34). Focal loss is designed to more highly weight pixel classifications that the model is ‘uncertain’ about, allowing the training to focus on sparse examples over easily classified background pixels. This is achieved by scaling standard cross entropy by a factor of (1 −*p*)^*y*^, where *p* is the model-estimated probability of the pixel belonging to the ground truth class, and *y* is a tuneable hyperparameter (35).

### 2.5. Training, Testing and Validation Data

A training library of paired images and manual labels was compiled from three-dimensional image data acquired both in-house and externally. Images from four modalities were chosen to represent a range of sources of contrast, imaging resolutions and tissues types (*Training 1-4*). As such, the training set encompasses a huge amount of variability with relatively little data. Volumes representing 25 % of both *Training 1* and *Training 2* were held out for model evaluation during hyperparameter tuning and training. An additional 3D MRI dataset (*Test 1*) provided unseen test data for model evaluation during training. Finally, additional data from modalities not included in the training/testing libraries were used to validate the cross-modality generalisability of the model (*Validation 1-3*). A summary of all datasets, with image sizes in pixels, is given in table 2. Details of how the images were generated are given below:

- *Training 1*: A high resolution episcopic microscopy (HREM) dataset, acquired in house by staining a healthy mouse liver with Eosin B and imaged using a standard HREM protocol (36).
- *Training 2*: X-ray microCT images of a microvascular cast, taken from a subcutaneous mouse model of colorectal cancer (acquired in house).
- *Training 3*: Raster-Scanning Optoacoustic Mesoscopy (RSOM) data from a subcutaneous tumour model (provided by Emma Brown, Bohndiek Group, University of Cambridge).
- *Training 4*: retinal angiography data obtained using Optical Coherence Tomography Angiography (OCT-A) (provided by Dr Ranjan Rajendram, Moorfields Eye Hospital).
- *Test 1*: T1-weighted Balanced Turbo Field Echo Magnetic Resonance Imaging (MRI) data from a machine-perfused porcine liver, acquired in-house.
- *Validation 1*: a subcutaneous colorectal tumour mouse model was imaged in house using Multi-fluorescence HREM in house, with Dylight 647 conjugated lectin staining the vasculature (36).
- *Validation 2*: an MF-HREM image of the cortex of a mouse brain, stained with Dylight-647 conjugated lectin, was acquired in house (23).
- *Validation 3*: two-photon data of mouse olfactory bulb blood vessels, labelled with sulforhodamine 101, was kindly provided by Yuxin Zhang at the Sensory Circuits and Neurotechnology Lab, the Francis Crick Institute.

Manual labelling of blood vessels was carried out using Amira (2020.2, Thermo-Fisher, UK). Fig. 1 shows representative 2D sections from the training and test images alongside the manual labels.

**Table 1.** Abbreviations and parameter definitions.

| Abbreviation/parameter | Definition |
| --- | --- |
| *cl* | Centreline sensitivity |
| CT | Computed Tomography |
| FN | False negative |
| FP | False positive |
| *H* | Hausdorff distance |
| HiP-CT | Hierarchical Phase Contrast Microscopy |
| MR-HREM | Multi-fluorescent High Resolution Episcopic Microscopy |
| MRI | Magnetic resonance imaging |
| OPT | Optical projection tomography |

**Table 2:** Summary of data used in training and validation.

| Dataset | Modality | Tissue | Volume (pixels) | Volume (mm) |
| --- | --- | --- | --- | --- |
| Training 1 | Eosin HREM | mouse liver | $4080 \times 3072 \times 416$ | $9.2 \times 6.9 \times 0.72$ |
| Training 2 | CT | mouse subcutaneous tumour | $1000 \times 1000 \times 682$ | $20 \times 20 \times 13$ |
| Training 3 | RSOM | mouse subcutaneous tumour | $191 \times 221 \times 400$ | $3.8 \times 4.4$ |
| Training 4 | OCT-A | human retina | $500 \times 500 \times 64$ | $12 \times 12 \times 1.5$ |
| Test 1 | MRI | porcine liver | $400 \times 400 \times 15$ | $360 \times 360 \times 75$ |
| Validation 1 | MF-HREM | mouse subcutaneous tumour | $2659 \times 2034 \times 1561$,<br>from which a subvolume of<br>$480 \times 480 \times 480$ was<br>used in fine-tuning | $9.2 \times 6.9 \times 2.7$ |
| Validation 2 | MF-HREM | mouse cortex | $4080 \times 3072 \times 99$ | $2.3 \times 1.75 \times 0.17$ |
| Validation 3 | Two photon microscopy | mouse olfactory bulb | $1536 \times 1536 \times 79$,<br>from which a subvolume of<br>$500 \times 250 \times 79$ was<br>used in fine-tuning | $0.3 \times 0.7 \times 0.4$ |

### 2.6. Training

Training data were prepared by normalising pixel intensities between zero and one. Optionally, the data was cropped at this point, for example in datasets where only a small region was of interest. Pixel data and true labels were converted into numpy arrays and saved in numpy format (npy) files. This file type is optimised for faster loading and occupies less memory than alternative formats, but lacks a metadata header. Therefore, an additional header class was created for easier data curation (Fig. 1).

Due to the large volume of data used in training, memory constraints became a limiting factor. Therefore, instead of loading entire data sets into motherboard RAM for training, a mini-batch of 64 × 64 × 64 pixel sub-volumes were read in from the disk at the start of each iteration (Fig. 1). This method of loading in pixel data increases the time taken for training but allows the model to be trained on more data without running out of memory. It also has the additional effect of acting as a translation augmentation of the data, as the sub-volumes were taken from random sets of coordinates within the training data. Additional augmentations are then performed on the generated subvolumes, randomly selecting from the following: rotations in the xy plane between −30^°^ and 30^°^; rescaling by a factor between 1.0 and 1.25; reflection in one or both of the in-plane axes.

A range of optimisers and learning rate schemes were compared to improve speed of convergence and final accuracy of the model, with the Adam optimiser (37) and an exponential decay learning rate schedule ultimately being chosen for training. The model was trained with a batch size of six, which was the maximum that could be stored in the VRAM of the GPUs used for training, and stopped when no further improvement was seen in the accuracy of the validation set. Training took approximately 24 hours.

### 2.7. Evaluation

Model performance was measured by calculating the area under the Receiver Operating Characteristic (AUROC) curve, Average Precision (AP) and DICE score. Precision and recall were calculated voxel-wise. The ability of the model to generalise to new data was assessed by calculating AUROC, AP and DICE scores for manually labelled sub-volumes of Validation images 1, 2 and 3 (in the case of Validation 1 and 3, where fine tuning was performed, a manually labelled sub-volume was withheld for evaluation).

## 3. Results

### 3.1. Accounting For Class Imbalance

Three loss functions designed to address imbalanced classes were compared in order to identify the best loss function for the tUbenet model. Training with the weighted categorical crossentropy (WCCE) loss function produced the most accurate segmentation, achieving an AUROC of 0.9 on the validation data (Fig. 2), but required iterative hyperparameter optimisation to identify the best combination of weights (in this case, 1:7). DICE-Binary crossentropy (DICE-BCE) outperformed focal loss (AUROC of 0.86 versus 0.80) and had the advantage of not requiring hyperparameter tuning. As such, DICE-BCE is used throughout this work unless otherwise stated.

**Figure 2:**
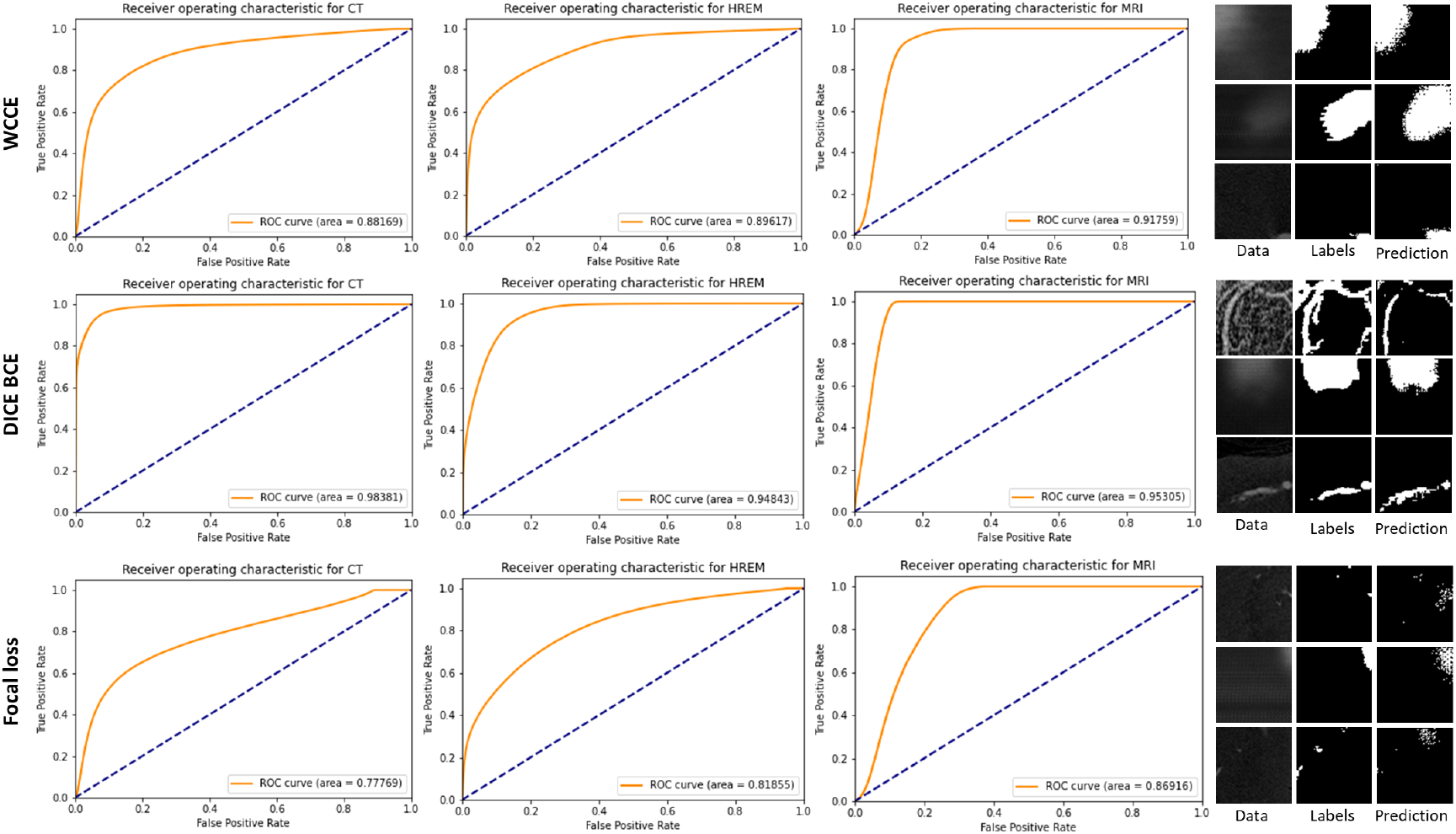
Comparison of ROC curves for three different loss functions (WCCE, DICE-BCE and focal loss), for two validation datasets. The left-hand panel shows the central slice of example 64 × 64 × 64 volumes, along with the ground truth and predicted labels for each model. DICE BCE outperformed both the WCCE and focal loss functions, with focal loss producing the lowest AUROC score and visibly worse segmentations.

### 3.2. Cross-branch Attention

The inclusion of a cross-branch attention mechanism sped up the initial rate at which the model learned the segmentation, with a AUROC score of 0.9 being achieved within 60 epochs, versus 200 epochs for the model without attention (Fig. 3). However, the attention model did not improve beyond 60 epochs, and both model versions converged at approximately the same level of accuracy. This result differs from the marked improvement in segmentation accuracy seen in Ref. 38. Despite converging in fewer epochs, the attention model had a much higher computational cost and a slower inference step. For these reasons the standard model structure without attention was preferred, and will be used throughout the rest of this work.

**Figure 3:**
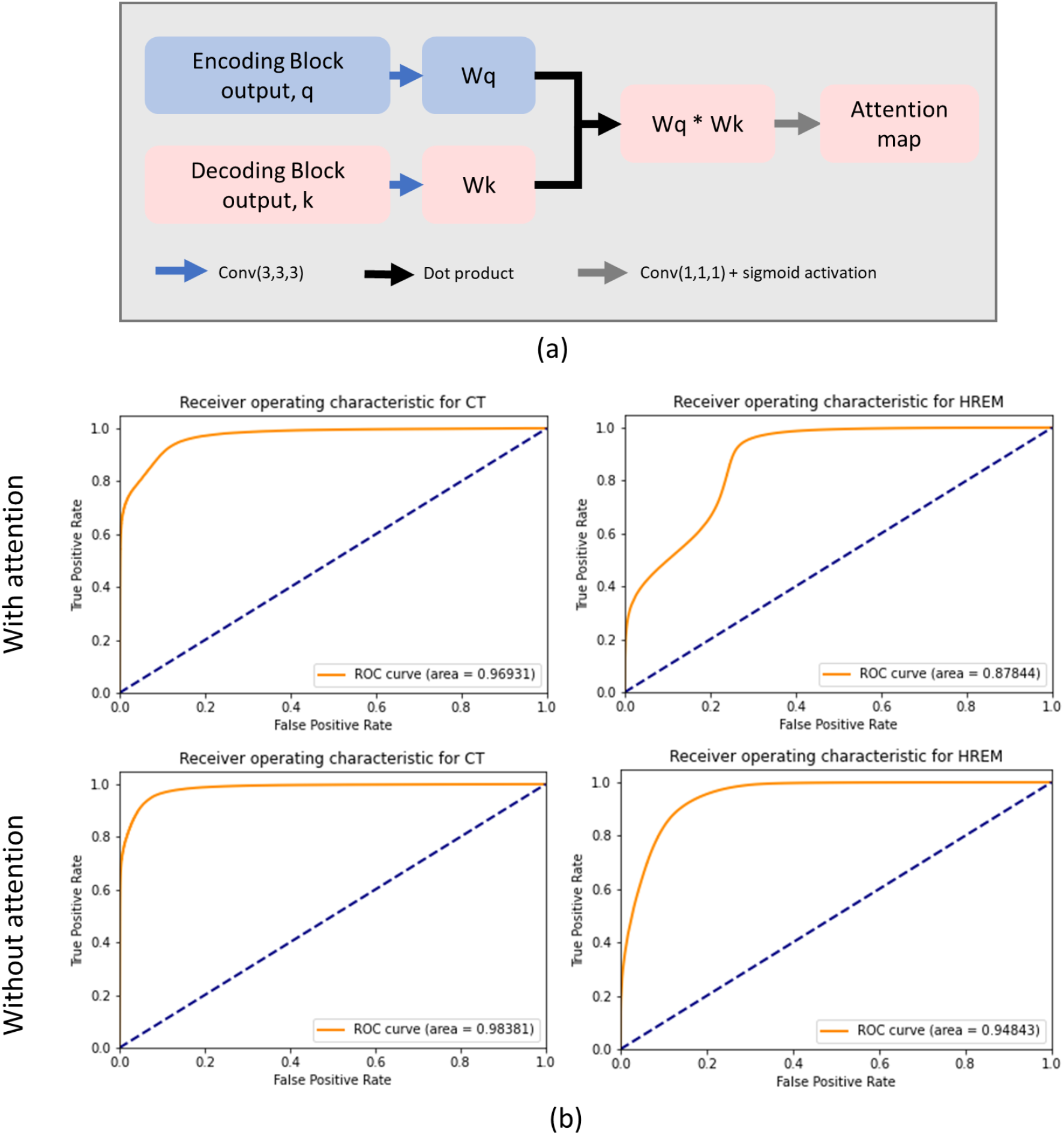
(a) A schematic illustration of the attention method used in this model. Concatenation operations, represented by the grey arrows in Fig. 1, were replaced with an attention layer as shown here. The feature maps from corresponding encoding and decoding layers (q and k respectively) are convolved with trainable kernels to produce Wq and Wk. These outputs are combined to find the dot product. A final convolution and sigmoid activation function produces the attention map for this layer, which is multiplied by the decoding feature map. (b) Receiver operating characteristic curves for the model with (top) and without (bottom) the inclusion of attention layers. The inclusion of attention did not improve segmentation quality when tested on the two validation datasets.

### 3.3. Cross-modality Generalisation

Through training on a library of data from different imaging modalities, the aim was to create a vessel-labelling model that would generalise to any volumetric dataset, even those acquired by previously unseen modalities, with minimal fine tuning. To test whether this aim had been achieved, tUbenet was evaluated on positive-contract MF-HREM images of a subcutaneous hypopharyngeal cancer model (36). This modality was chosen due to the relative newness of MF-HREM: there is very little labelled data available for training, meaning existing deep learning approaches would not be suitable for analysing this data.

Applying the trained model without fine-tuning to this new data type, we produced a segmentation of the vasculature with high recall of the vessel class (96 %) but low precision (23 %) (2 s.f.). Erroneous labelling of out-of-plane fluorescence as vasculature was noted, leading to the low precision score (Fig. 4). However, after two hours of fine-tuning the model using a labelled subvolume of the data (120 × 120 × 120 pixels) representing just 0.3 % of the total image volume, the recall and precision were improved to 0.98 and 0.97 respectively (DICE score = 0.97, AP = 0.87) (2 s.f.) (Table 3). Increasing the train set size to 480 × 480 × 480 pixels (1.2 % of the total image volume) further improved the segmentation accuracy (precision 0.99, recall 0.97, DICE 0.98, AP 0.97 (2 s.f.)). This sub-volume was manually annotated in a matter of hours, compared to the days/weeks that would be required to label the entire dataset. This result demonstrates that our pre-trained model can be fine-tuned to new data with a minimal amount of manually annotation, making it a practical option for applications where large libraries of labelled data do not exist.

**Table 3:** Effect of data subvolume size used in fine-tuning.

| Training Volume | DICE Score | AP |
| --- | --- | --- |
| $120 \times 120 \times 120$ | 0.97 | 0.87 |
| $240 \times 240 \times 240$ | 0.98 | 0.94 |
| $480 \times 480 \times 480$ | 0.98 | 0.97 |

**Figure 4:**
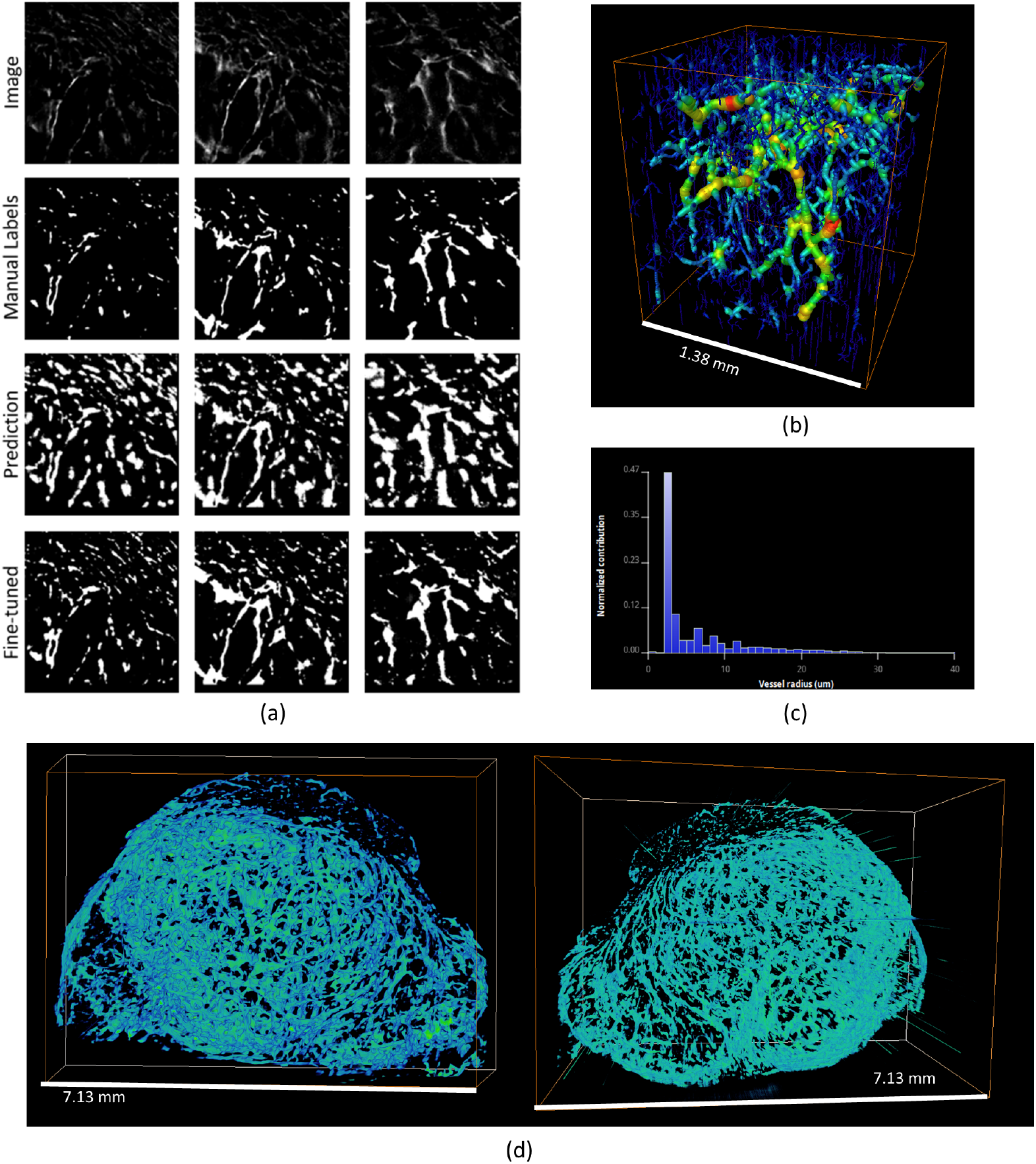
(a) 2D sections from an MF-HREM dataset of a fluorescence-stained subcutaneous tumour (top row) are shown alongside the corresponding manual labels (second row) and the predicted labels generated by the pre-trained model prior to fine-tuning (third row) and following fine-tuning (bottom row). (b) A skeletonised subvolume of the tumour and corresponding vessel radii distribution (c). (d) 3D views of the entire dataset segmented using the fine-tuned model.

Moreover, the fine-tuned model was then applied to another MF-HREM dataset from a different tissue: the cortex of a mouse brain stained with DyLight-647 conjugated lectin (Fig. 5). Despite differences in cortical and tumour vasculature, the model was able to generalise to the cortex dataset with a high level of accuracy and no additional fine-tuning (recall=0.90, precision=0.88, DICE score=0.89 (2 s.f.)). This suggests that fine-tuning may only need to be done once per image type, further saving on manual labelling.

**Figure 5:**
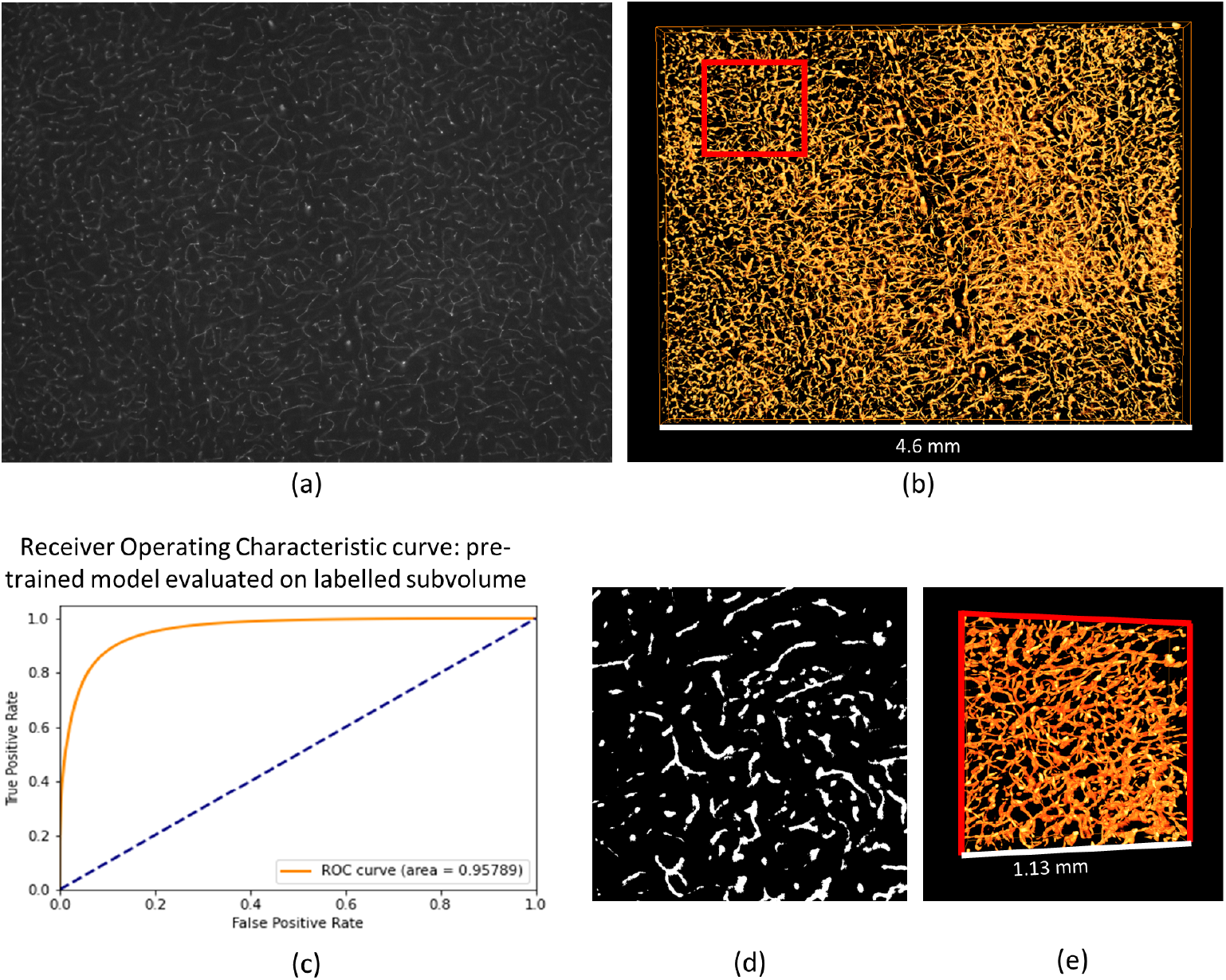
(a) 2D section of an MF-HREM image of the cortical vasculature of a mouse. (b) 3D vessel segmentation produced by the tubenet model fine-tuned using the tumour MF-HREM dataset seen in Fig 4. The red square highlights a region of the data that was manually labelled to provide a ground truth for validation. (c) ROC curve analysis performed using the manually labelled ground truth (d) and model predictions for this subvolume (e) shows that the prediction is highly accuracy (AUROC = 0.96 (2 s.f.)).

Finally, the model was applied to images of the blood vessels of the mouse olfactory bulb, acquired by two photon microscopy. This dataset provided an additional challenge as, not only it an imaging modality that the model had not previously seen, but it exhibits spatially varying imaging artefacts due to resolution drop off with depth. In order to specialise the model to this data, a sub-volume that spanned the whole z-axis (79 virtual sections) was manually labelled for fine-tuning. Training on a volume of 79 × 500 × 250 pixels (5 % of the whole image volume) resulted in a model that labelled vasculature with a precision of 0.84 and recall of 0.98 (2 s.f.) (Fig. 6).

**Figure 6:**
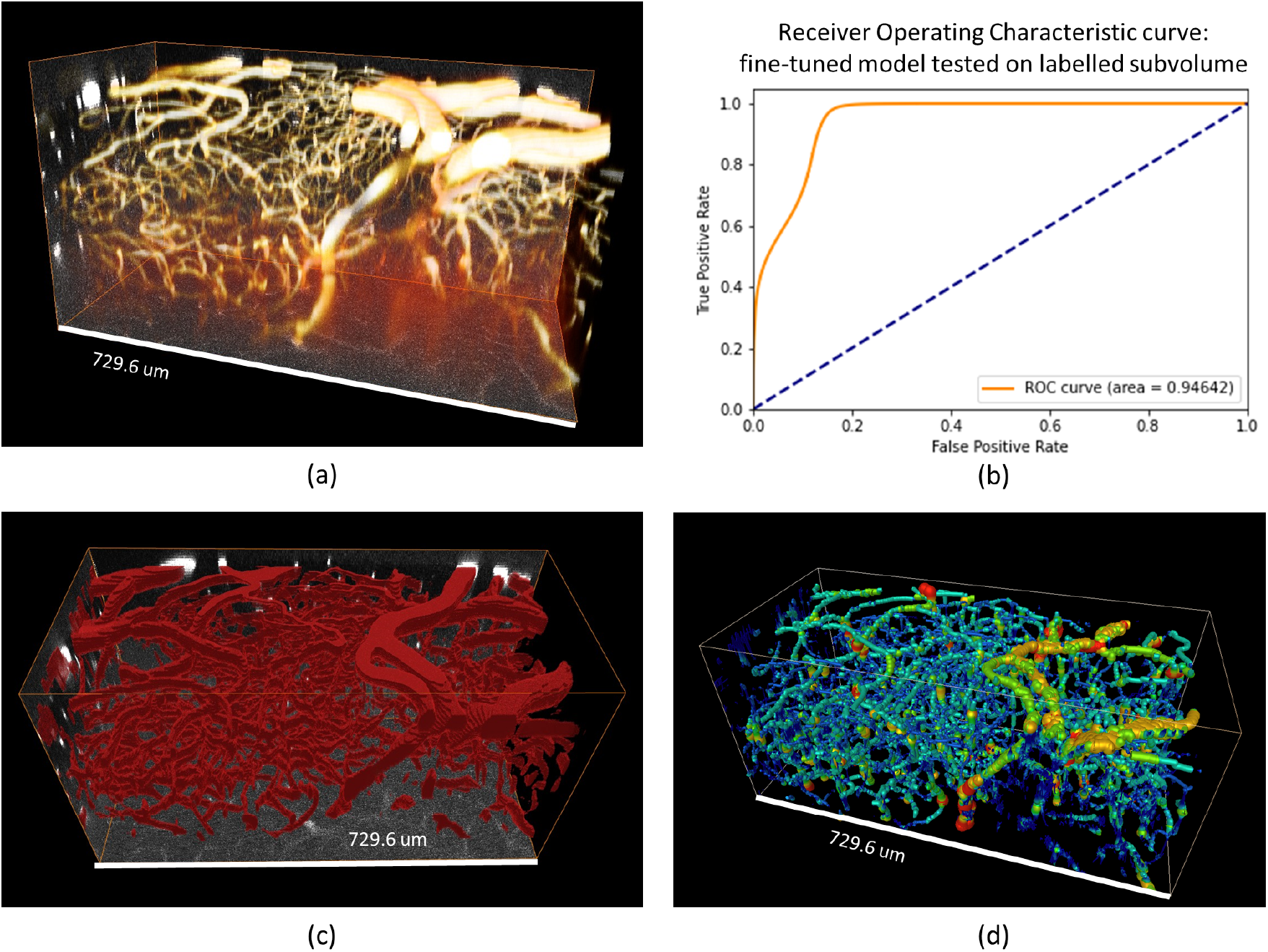
(a) Two photon fluorescence microscopy data showing labelled vasculature (yellow) in the mouse olfactory bulb. (b) ROC curve analysis of the model prediction following fine-tuning on a manually labelled subvolume of the data shows that the model is accurately labelling vasculature (AUROC = 0.95 (2 s.f.)). (c) A 3D visualisation of the predicted vessel labels (red). (d) Skeletonised vasculature, colour coded by vessel radius.

## 4. Discussion

In this work a 3D U-net for vessel segmentation was trained on a highly varied dataset, comprised of four imaging modalities and three tissue types. The general model was then specialised for new applications - segmenting data from previously unseen modalities - with minimal fine-tuning. Our results show that this two-step training approach to deep learning for image analysis is a viable strategy. Fine-tuning the general model with a small sub-volume of the new data of interest (as little as < 1% of the data volume) was sufficient to generate accurate label predictions for the entire dataset. This approach to deep learning is especially advantageous when analysing data from new or lesser-used imaging modalities, where a back catalogue of data is not available to train a model from scratch.

In the future this model could be made more accessible to non-computational researchers creating a complementary graphical user interface for manual labelling and fine-tuning, or by integrating it into existing open-source data visualisation programs (for example, as a Napari plug-in). Overall, the move towards an open-source, generalisable DL tool for vessel segmentation expands the options available to researchers looking to quantify vaculature in 3D and overcomes many of the bottlenecks preventing wider use of DL in the 3D imaging community.

## Acknowledgments

Thank you to Dr Ranjan Rajendram (Moorfields Eye Hospital), Yuxin Zhang and Dr Carles Bosch Piñol (Francis Crick Institute), Dr Emma Brown and Dr Sarah Bohndiek (University of Cambridge) for kindly providing data that was used in model training and validation.

Furthermore, thank you to Tian Ding, Monica Sidarious and James Gledhill for contributing to the manual segmentation of the training data.

This research was funded by Cancer Research UK (C44767/A29458 and C23017/A27935) and EPSRC (EP/W007096/1).

Dr Walsh was supported by an MRC Skills Development fellowship (MR/S007687/1).

